# Experimental Validation of Coiled-Coil Architecture and Folding Dynamics in the Golgin Bug1

**DOI:** 10.64898/2026.09.21.753216

**Authors:** Marilia D.O. Silva, Raphael V.R. Dias, Igor S. Pedriz, Italo A. Cavini, Mayra T.S. Silva, Rodrigo J.S. Sena, Humberto D’Muniz Pereira, Fernando A. Melo, Luis F. S. Mendes

## Abstract

Golgins are widely described as long coiled-coil proteins that contribute to the structural organisation and trafficking functions of the Golgi apparatus. Although experimental structures have been determined for a limited number of golgin regions, atomic-level information on their extended coiled-coil segments remains scarce, and the oligomeric state, topology, and register of most predicted regions remain unestablished. Here, we characterise a predicted coiled-coil region of the yeast golgin Bug1 (BUG1cc) using structural, biophysical, and computational approaches. X-ray crystallography revealed a parallel, in-register dimeric coiled-coil containing ten heptad repeats and a predominantly hydrophobic core, with specific polar interactions contributing to dimer stabilisation. In solution, BUG1cc was dimeric under SEC-MALS conditions and remained highly α-helical across the pH and ionic strength conditions examined. CD measurements revealed pronounced scan-rate-dependent hysteresis, while DSC independently confirmed an asymmetry between heating and cooling transitions. Increasing protein concentration shifted both apparent transition temperatures while preserving thermal hysteresis, supporting chain association and conformational rearrangements in structural recovery. Structure-based simulations indicated that interface contacts and intra-chain helicity are thermodynamically coupled and melt as a single cooperative unit, and that the monomer released on dissociation is compact and only partially helical, so that reassociation proceeds through a coupled folding-binding mechanism whose rate-limiting step is conformational rather than bimolecular. Together, these results establish the molecular architecture of a predicted coiled-coil region of Bug1 and reveal a complex folding landscape in which oligomerisation, secondary-structure recovery, and kinetic barriers are tightly coupled.

## Introduction

The Golgi apparatus is a central organising station of the eukaryotic secretory pathway, where proteins and lipids are modified, sorted, and redistributed with spatial and temporal precision. Golgi morphology directly conditions trafficking efficiency, glycosylation fidelity, and the biochemical environment through which cargoes mature *en route* to the cell surface, the endolysosomal system, or the extracellular space (*1*). Recent work has reinforced this view by showing that Golgi organisation is functionally coupled to the distribution of glycosylation enzymes, cargo processing, and the maintenance of distinct secretory outputs, emphasising that Golgi structure determines cell physiology rather than passively reflecting membrane traffic (*2, 3*).

This architectural control has long motivated interest in the so-called Golgi matrix, a set of Golgi-associated proteins mainly composed of GRASPs and golgins (*4*). These protein families are considered key to Golgi stacking, ribbon organisation, cisternal identity, and vesicle tethering (*5–8*). Recent evolutionary and cell-biological analyses have further expanded their relevance. Comparative work has suggested that coordinated interactions between GRASPs and golgins were important in the emergence of ribbon-like Golgi organisation in animals. At the same time, recent functional studies indicate that individual golgins can shape the physical and biochemical space required for efficient secretion and extracellular matrix assembly (*9, 10*). Together, these findings reinforce the notion that Golgi matrix proteins actively organise Golgi form and function.

Within this context, golgins occupy a particularly prominent position (*11*). They are generally considered long-range tethering factors that extend from Golgi membranes to capture incoming transport carriers, thereby helping to impose directionality and specificity on membrane traffic (*5, 12, 13*). This view remains central in current models of Golgi function and has been refined by studies showing that different golgins capture distinct classes of vesicles and contribute nonredundantly to Golgi homeostasis (*14*). Recent literature has also begun to move beyond the classical picture of golgins as isolated tethering rods. In this context, GM130 was recently shown to coordinate with RNAs to scaffold the Golgi ribbon through a protein–RNA condensation mechanism driven by phase separation (*7*). A 2025 preprint from Rothman’s group, based on *in situ* super-resolution microscopy and biochemical reconstitution, proposed that multiple rim golgins are organised in a layered tetraplex at the Golgi edge and that purified golgins can self-assemble into filamentous bands (*6*). Although these observations still await the full weight of peer review, they underscore a growing shift in the field: golgins are increasingly discussed not only as vesicle tethers, but also as higher-order structural elements of Golgi organisation.

A defining feature of golgins is their high content of predicted coiled-coil regions, which has shaped current models of their elongation, oligomerisation, and long-range tethering activity (*12, 15, 16*). Experimental structural information is available for a limited number of golgin domains and short coiled-coil-containing regions, including the coiled-coil-containing Rab6-binding region of GCC185, which was structurally determined in complex with Rab6 (*17*). Nevertheless, most extended coiled-coil segments that constitute golgin scaffolds remain structurally uncharacterised. Consequently, their oligomeric state, parallel or antiparallel topology, helical register, and residue-specific stabilising interactions are generally inferred from sequence predictions rather than determined experimentally. This distinction matters because coiled-coil predictions, while informative, do not uniquely define these structural properties and cannot exclude alternative oligomeric arrangements (*18*).

This issue is particularly relevant for fungal Golgi proteins, where family-level structural assumptions have often travelled faster than direct molecular characterisation. In budding yeast, Bug1 (Ydl099w; UniProt Q12191) was identified as a Golgi-associated partner of the GRASP ortholog Grh1 and functionally linked to ER-to-Golgi traffic through genetic and biochemical interactions with key tethering factors, including Uso1 and Ypt1 (*19, 20*). Since its initial description, Bug1 has been referred to as a coiled-coil protein and has been annotated accordingly (*20*). Yet the molecular architecture of Bug1 has not been experimentally determined, and its classification as a coiled-coil protein rests primarily on sequence-based prediction and functional context. In particular, its oligomeric state, topology, register, and atomic interactions in the predicted coiled-coil region remain unknown.

Bug1 therefore provides a useful model for determining how a predicted golgin coiled-coil region is organised at the molecular level. Here, we combine X-ray crystallography, solution biophysics, and structure-based simulations to investigate residues 185–273 of Bug1. We show that this segment forms a parallel, in-register homodimer containing ten heptad repeats and remains dimeric and predominantly α-helical in solution. Thermal measurements further reveal scan-rate-dependent hysteresis, indicating a substantial kinetic barrier to recovery of the native assembly. These results establish the atomic architecture of the Bug1 coiled-coil domain and provide a framework for investigating golgin coiled-coil folding and assembly.

## Materials and Methods

### Protein expression and purification

The region encoding residues 185–273 of Saccharomyces cerevisiae Bug1 (UniProt Q12191) was predicted using coiled-coil predictors through Waggawagga (21), amplified by PCR, and cloned into a pET-SUMO expression vector. A tryptophan residue was introduced immediately upstream of Bug1 residue S185 to enable protein quantification by absorbance at 280 nm. After ULP1 cleavage of the SUMO tag, an additional serine residue remained at the N-terminus. The final recombinant BUG1cc construct therefore contained 91 residues, comprising the 89-residue native Bug1 segment and two additional N-terminal residues, Ser and Trp. Its final sequence was: SWSDFLTTIKKQKEEDELTKLRAENEKLTQENKQLKFLNMENETTVDDLQDQLQEKEDIINGLQNDLQTARDELIAAVEKLKLAEAKAARN.

The resulting construct was verified by DNA sequencing and transformed into Escherichia coli Rosetta2 (DE3) cells. Protein expression was induced at an optical density at 600 nm of 0.6–0.7 by addition of 0.5 mM IPTG, followed by incubation for 16 h at 18 °C. Cells were harvested by centrifugation and resuspended in 20 mM Tris-HCl, pH 7.5, containing 500 mM NaCl and 40 mM imidazole. Following sonication, the lysate was clarified by centrifugation at 25,000 × g for 25 min. The supernatant was loaded onto a cobalt-affinity resin equilibrated in the same buffer. After washing, the SUMO-tagged protein was eluted with 20 mM Tris-HCl, pH 7.5, containing 500 mM NaCl and 500 mM imidazole. The SUMO tag was removed by overnight incubation with ULP1 during dialysis against 20 mM Tris-HCl, pH 7.5, containing 500 mM NaCl. The sample was subsequently reapplied to the affinity resin, and untagged BUG1cc was recovered in the flow-through. Final purification was performed by size-exclusion chromatography using a Superdex 75 10/300 GL column equilibrated in Buffer A for crystallographic trials (20 mM Tris-HCl, 50 mM NaCl, pH 7.5) and Buffer B for thermodynamic experiments (10 mM Sodium Phosphate, 50 mM NaCl, pH 7.5) at a flow rate of 0.5 mL/min. Protein purity was assessed by SDS-PAGE.

### Circular dichroism spectroscopy

Circular dichroism (CD) measurements were performed using a J-815 spectropolarimeter (Jasco) equipped with a Peltier temperature-control system. Far-UV CD spectra were recorded in a 1 mm path-length quartz cuvette from 190 to 270 nm using a scan speed of 50 nm min⁻¹, a spectral bandwidth of 1 nm, and a digital integration time of 2 s. Unless otherwise indicated, BUG1cc samples were diluted to 0.1 mg mL⁻¹ in 20 mM sodium phosphate buffer, pH 7.4. This represented at least a 20-fold dilution of the protein stock, minimising differences in buffer composition among samples. The corresponding buffer baseline was subtracted from each spectrum. CD spectra were recorded in three independent measurements and deconvoluted using BeStSel (22).

Temperature-dependent far-UV CD spectra were collected between 20 and 70 °C at 5 °C intervals to monitor changes in secondary structure. Thermal unfolding and refolding were monitored independently at 222 nm during heating and cooling between 10 and 70 °C. Heating and cooling scans were conducted at 0.5, 1, 2, 5, 7, and 10 °C min⁻¹, with three replicate measurements performed at each scan rate. The apparent transition midpoint for each branch, T_M_, was determined by the midpoint of a sigmoidal curve fit, and thermal hysteresis was calculated as Δ*T_M,app_* = (*T_M,ℎeating_* − *T_M,cooling_*).

The effect of pH was examined using BUG1cc samples prepared in 10 mM sodium phosphate buffer adjusted to pH 5.5, 6.5, 7.0, 7.5, or 8.0. The effect of ionic strength was evaluated in 10 mM sodium phosphate buffer (pH 7.5) containing 50 or 150 mM NaCl. A nominally low-salt condition was prepared without the addition of NaCl; however, because the protein stock contained NaCl, this condition still had an estimated final NaCl concentration below 10 mM after dilution. The θ_222_/θ_208_ ratio was calculated from the corresponding baseline-corrected CD intensities.

The effect of protein concentration on thermal unfolding and refolding was examined using BUG1cc at 0.02, 0.05, 0.1, 0.2, 0.4, and 0.6 mg mL⁻¹, with heating and cooling rates fixed at 5 °C min⁻¹. We performed four independent measurements at each concentration. For this series, apparent transition temperatures (T_m,app_) were determined from the dominant peak in the magnitude of the first derivative of the CD signal at 222 nm with respect to temperature, or by a single sigmoid Boltzmann fit when applicable. We also fitted normalised cooling profiles at 0.4 and 0.6 mg mL⁻¹ with single and double Boltzmann sigmoid functions with constant baselines to characterise asymmetry. The double-sigmoid function comprised the sum of two sigmoidal components with independently fitted centres and widths and a fitted relative amplitude. Data are reported as mean ± standard deviation from four independent measurements.

### Differential scanning calorimetry

Differential scanning calorimetry (DSC) measurements were performed using a VP-DSC MicroCal calorimeter (MicroCal, Northampton, MA, USA). BUG1cc was prepared at 3 mg/mL in 10 mM phosphate buffer (pH 7.5) containing 50 mM NaCl. Thermograms were recorded from 5 to 70 °C at a scan rate of 90 °C/h. After reaching the upper temperature, samples were held for 10 min and subsequently cooled at the same rate. Raw thermograms were baseline-corrected by subtracting the corresponding buffer trace and subsequently normalised to the molar concentration of BUG1cc chains, calculated using a molecular mass of 11 kDa. Accordingly, ΔH_cal_ values are reported per mole of BUG1cc chain. Thermal baselines (cubic, progressive, or manual) were reconstructed and subtracted from the normalised traces using MicroCal Origin software. The apparent van’t Hoff enthalpy was estimated from the full width of the calorimetric transition at half maximum according to

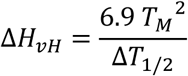

where T_M_ is the temperature at the maximum excess heat capacity, expressed in kelvin, and ΔT_1/2_ is the full width of the transition at half maximum, also expressed in kelvin. The resulting ΔH_vH_ values were expressed in cal.mol⁻¹ and subsequently converted to kcal.mol⁻¹. The calorimetric enthalpy, ΔH_cal_, was obtained by integration of the baseline-corrected excess heat-capacity peak. Heating and cooling thermograms were analysed independently. Because the transitions exhibited hysteresis, T_M_, ΔH_vH_, and ΔH_cal_ were treated as apparent parameters determined at the applied scan rate rather than as equilibrium thermodynamic quantities. Each experiment was independently repeated twice to ensure reproducibility of the data.

### Size-exclusion chromatography coupled to multi-angle light scattering (SEC-MALS)

Size-exclusion chromatography coupled with multi-angle light scattering (SEC-MALS) was performed using a miniDAWN TREOS detector (Wyatt Technology, CA) equipped with three detection angles (43.6°, 90°, and 136.4°) and a 659 nm laser. A Wyatt QELS module was used in-line for dynamic light scattering (DLS) measurements to determine the hydrodynamic radius (R_H_), along with an Optilab® rEX refractive index detector (Wyatt Technology). All measurements were carried out using a Superdex 200 HR 10/300 analytical SEC column (GE Healthcare). Bovine serum albumin (BSA; Sigma-Aldrich) was used as a reference standard. Protein samples (∼2 mg/mL) were eluted in 20 mM Tris-HCl, 50 mM NaCl, pH 8.0, at a constant flow rate of 0.5 mL/min. Immediately before data acquisition, the samples were centrifuged at 10,000 × g for 10 minutes at 4°C to remove aggregates or debris. Data acquisition and analysis were performed using ASTRA 7 software (Wyatt Technology) with the following parameters: refractive index of 1.331, solvent viscosity of 0.890 cP, and a refractive index increment (dn/dc) of 0.185 mL/g. SEC-MALS data were measured in 2 independent measurements.

### Crystallisation and X-ray diffraction data collection

Initial crystallisation screening was performed using the sitting-drop vapour-diffusion method with commercial screening kits and a crystallisation robot. Promising conditions were identified in solutions containing 0.1 M HEPES (pH 7.0), 10% (w/v) PEG MME 5000, and 5% Tacsimate. These conditions were subsequently optimised manually by systematically varying the PEG MME 5000 concentration (6, 8, 10, 12, 14, and 16% [w/v]) and pH (6.6, 6.8, 7.0, 7.2, 7.4, and 7.6), using solutions based on 0.1 M HEPES, PEG MME 5000, and 5% (v/v) Tacsimate. Multiple crystals were obtained under the optimised conditions, three of which exhibited diffraction quality suitable for X-ray structural determination. The crystal used for structural determination was grown in 0.1 M HEPES pH 7.2, 14% PEG MME 5000, and 5% (v/v) Tacsimate

### X-ray data collection, structure determination and refinement

X-ray data were collected at Sirius MANACÁ, CNPEM, Brazil. Processing was performed using XDS via AutoProc. Anisotropic correction of the data was performed using STARANISO 2.4.19 (Global Phasing) (https://staraniso.globalphasing.org/cgi-bin/staraniso.cgi, accessed in 2026). Ab initio phasing was done by using 21-aa polyalanine helices with ARCIMBOLDO LITE (*23*) in coiled-coil mode. Model refinement was performed using Buster 2.10.3 (Global Phasing). The helical register was determined by manually inspecting the electron density of side-chain core residues. The three-dimensional model was refined by iterative manual building into 2*F*_o_-*F*_c_ and *F*_o_-*F*_c_ electron density maps using Coot (*24*). Torsion/libration/screw (TLS) motion restraints were turned on throughout the final refinement steps. Model validation was done using MolProbity (*25*).

**Table 1.** Data collection and processing. Values in parentheses correspond to the outer shell.

|  |  |
| --- | --- |
| Diffraction source | Sirius MANACÁ |
| Wavelength (Å) | 0.9772 |
| Detector | Pilatus 2M |
| Crystal-detector distance (mm) | 175.87 |
| Rotation range per image (°) | 0.1 |
| Total rotation range (°) | 360 |
| Space group | $C_2$ |
| $a, b, c$ (Å) | 110.44, 42.45, 63.35 |
| $\alpha, \beta, \gamma$ (°) | 90.00, 110.25, 90.00 |
| Anisotropy analysis |  |
| Diffraction limits 1, 2, 3 (Å) | 1.682, 2.173, 2.140 |
| Direction 1<sup>†</sup> | 0.9433, 0.0000, 0.3319; $a^* - 0.009 c^*$ |
| Direction 2<sup>†</sup> | 0.0000, 1.0000, 0.0000; $b^*$ |
| Direction 3<sup>†</sup> | -0.3319, 0.0000, 0.9433; $-0.501 a^* + 0.866 c^*$ |
| Resolution range (Å) | 48.17-1.86 (1.97-1.86) |
| Total No. of reflections | 157915 (26050) |
| No. of unique reflections | 23521 (3718) |
| Completeness, spherical (%) | 72.7 (27.7) |
| Completeness, ellipsoidal (%) | 91.5 (79.2) |
| Multiplicity | 6.7 (7.0) |
| $\langle I/\sigma(I) \rangle$ | 1.893<sup>#</sup> (at 1.86 Å) |
| $CC_{1/2}$ | 0.996 (0.832) |
| $R_{\text{r.i.m.}}$ | 0.103 (0.352) |
| Reflections used in refinement | 17101 |
| $R, R_{\text{free}}$ | 0.249, 0.310 |
| Overall $B$ factor from Wilson plot (Å<sup>2</sup>) | 31.75 |
| Ramachandran favored (%) | 100 |
| Ramachandran allowed (%) | 0 |
| All-atom clashscore | 0 |
| Bond lengths (RMSD) (Å) | 0.008 |
| Bond angles (RMSD) (°) | 0.91 |
| PDB entry | 38EE |

<sup>†</sup>Directions of principal axes of the ellipsoid expressed as direction cosines in the orthogonal basis.

### System and structure-based model

The dimerization thermodynamics and folding mechanism of BUG1cc, a homodimeric coiled coil of two identical 91-residue chains (182 Cα beads in the model), were probed with Cα structure-based (native-centric, “Gō-like”) models (*26, 27*), in which each residue is represented by a single bead centered on its Cα atom and the energy function is defined so that the experimentally determined crystal structure sits at the global minimum of a minimally frustrated, funneled landscape. Because non-native interactions are absent by construction, any non-two-state behavior, intermediate, or kinetic bottleneck that emerges reflects native topology and chain connectivity rather than sequence-specific mispairing, the key distinction for a coiled coil whose cooperative character is under experimental scrutiny. Models were generated with SMOG 2 using the default Cα shadow contact map (non-bonded cutoff 1.1 nm) and simulated through the OpenSMOG/OpenMM interface (*28, 29*) on a GPU. Two systems were built from the same native template: the full dimer (both chains, inter- and intra-chain contacts) and the isolated monomer (one chain retaining only its intra-chain contacts), the latter providing the partner-free reference needed to separate intrinsic single-chain folding from bimolecular association effects.

### Replica-exchange simulations and free-energy reconstruction

Equilibrium thermodynamics were obtained by temperature replica-exchange molecular dynamics (REMD) (*30*) using 24 replicas spanning a non-uniform reduced-temperature ladder T* = 0.70–1.25, with spacing concentrated in the transition region (13 replicas at T* = 0.90–1.14) and sparser coverage in the native baseline (5 replicas, T* < 0.90) and the fully denatured plateau (6 replicas, T* > 1.14). The Langevin integrator used a time step DT = 0.0005 (in reduced units) and a friction coefficient γ = 1.0 (in inverse reduced time units). Exchange attempts were performed every 1000 steps; equilibrium observables (Q_inter_, Q_intra_, d_COM_, total energy) were recorded every 5000 steps; full-coordinate snapshots were saved every 50 000 steps. Each replica ran for 50 million steps, with the first 30% of each trajectory discarded as equilibration, yielding approximately 35,000 production frames per replica. Because the coiled coil dissociates and re-associates on timescales inaccessible to fixed-temperature runs, REMD is essential for this system. We applied a flat-bottom harmonic restraint on the inter-chain centre-of-mass distance (r_0_ = 5.0 nm, k = 10 kJ mol^−1^ nm^−2^) to the dimer simulations. This restraint is zero for d_COM_ ≤ 5.0 nm, covering all native and partially dissociated states and activates only beyond 5 nm to prevent irreversible monomer escape during high-temperature replicas, thereby enabling full dissociation–reassociation round trips within finite simulation time without perturbing the thermodynamically relevant states. We report temperature in reduced units T* and, where compared with experiment, map it to °C using a two-point affine calibration. The Q_inter_ midpoint was anchored to the experimental DSC heating T_M_ of 48.45 °C, and the scale was set so that the full width at half maximum of the simulated excess heat-capacity peak equals that of the DSC endotherm on heating (ΔT_1/2_ = 7.44 °C). Both widths were measured on baseline-subtracted profiles, the simulated baseline being obtained by linear interpolation between the pre- (T* < 0.80) and post-transition (T* > 1.18) regions; this gives a scale factor of 43.2 °C per unit T*, applied identically to every panel with a Celsius axis. Because the experiment fixes both the position and the width of the simulated transition, neither the midpoint nor the width is an independent prediction of the model, and the conclusions below are stated in reduced units wherever they do not depend on the calibration.

### Order parameters and free-energy analysis

We followed folding and association using the fraction of native contacts Q, defined smoothly as in Best, Hummer, and Eaton (32), with a threshold of 1.2 × r0 (r0 = native Cα–Cα distance for each contact pair). Two independent axes were maintained: Qinter (inter-chain interface contacts; dimer association) and Qintra (intra-chain contacts averaged over chains A and B). Because the native template contains no intra-chain contacts with sequence separation greater than eight residues, Q_intra_ reports exclusively on local helical structure, not tertiary packing. The two coordinates have very different denatured baselines (Q_inter_ falls to 0.01, Q_intra_ only to 0.42, reflecting the local contacts that persist in a disordered chain), so transition midpoints were defined as the temperature at which each coordinate reaches the mean of its own pre- and post-transition baselines, rather than at a fixed absolute value of Q; midpoints assigned in this way were cross-checked against C_v_ and against the susceptibility χ = N·Var(Q). Coupling between the two coordinates was quantified at fixed temperature by conditioning the ensemble on association state and comparing 〈Q_intra_〉 between associated (Q_inter_ > 0.6) and dissociated (Q_inter_ < 0.3) frames of the same replica, with uncertainties from the standard error of each conditional mean. Additional coordinates were the inter-chain centre-of-mass distance d_COM_, the radius of gyration R_g_, and the end-to-end distance Ree. Heat capacity Cv was obtained from WHAM energy fluctuations (C_v_ = Var(E)/(T*)2); transition midpoints were assigned from the inflection of the corresponding Q curve, cross-checked against Cv and the susceptibility χ = N·Var(Q). Two-dimensional free-energy surfaces F_(Qinter, Qintra)_ and F_(Rg, Ree)_ were constructed by WHAM reweighting at selected temperatures, and ensemble-averaged contact-frequency matrices were computed at three representative thermal states (native, transition, dissociated).

### Encounter and all-atom simulations

Non-equilibrium re-association dynamics were studied by coupled folding–binding encounter simulations: two monomers were initialised in compact, separated conformations (d_COM_ ≈ 4.0 nm, Ree ≈ 6.1 nm) and allowed to associate near the dimer T_M_ under the same Langevin integrator; the resulting encounter complex was then subjected to stepwise cooling (stabilisation branch) or heating (re-dissociation branch). We ran five independent replicates; one diverged numerically, with the chains escaping to unphysical separations, and was excluded from all analyses, leaving four. These trajectories establish whether the productive folding–binding route is sequential (fold-then-bind) or coupled (interface forms concurrently with chain extension) and document the directional asymmetry that underlies scan-rate-dependent hysteresis. As an independent structural consistency check, explicit-solvent all-atom MD simulations were run at 25, 37, 44, 50 and 60 °C (100 ns each, first 20 ns discarded as equilibration). These runs test whether the crystallographic dimer is structurally stable on the sub-microsecond timescale at each temperature; being short, unbiased, fixed-temperature simulations, they are not expected to sample dissociation or unfolding and were not used to extract transition temperatures.

## Results and Discussion

### BUG1cc adopts a parallel, in-register dimeric coiled-coil architecture

We solved the crystal structure of our Bug1 coiled-coil (CC) construct. The diffraction pattern is anisotropic (diffraction limits between 1.682 and 2.173 Å), and its correction, together with *ab initio* phasing using ARCIMBOLDO-LITE (*23*), enabled structural model determination. The structure is a dimeric, parallel, in-register CC and agrees well with its AlphaFold3 (AF3) prediction (*33*) (RMSD = 1.05 Å over 132 Cα atoms), apart from the termini and a few side-chain positions.

The N-terminus (S185-E199) does not form CC contacts, and S185-E196 in chain A unfurls to make contacts with a neighbouring chain, likely due to crystal packing (Figure 1A). Ten heptad repeats are present (Figure 1B) from L200 (position *a*) to A270 (position *a*), containing mainly hydrophobic residues in the core (16 out of 21); the exceptions are K238 at *d*, which positions the side-chain amine out of the core, and N207, N214, N221 (all at *a*) and N224 (at *d*). Asparagine residues are polar and, at core CC positions, help specify oligomeric state (at *a*, more prevalent in dimers; at *d*, in trimers) and partner interaction (*34*). All three Asn at *a* form symmetric inter-chain contacts with the corresponding residue in the opposing chain. N224 at *d* is stabilised by three hydrogen bonds involving its N^δ2^ and *i*) one of the O^ε^ from E225’, *ii*) O^δ1^ from N221’ (both interchain) and *iii*) carbonyl O from L220 (Figure 1C). The presence of these asparagines in the core and the specific interactions involving N224 likely contribute to the specificity and stability of the BUG1cc homodimeric interface, despite the presence of many other CC proteins in the cell. Notably, AF3 and other deep-learning modelling methods did not predict some of these contacts correctly, which emphasises the importance of experimental models for accurate atomic analysis.

**Figure 1.**
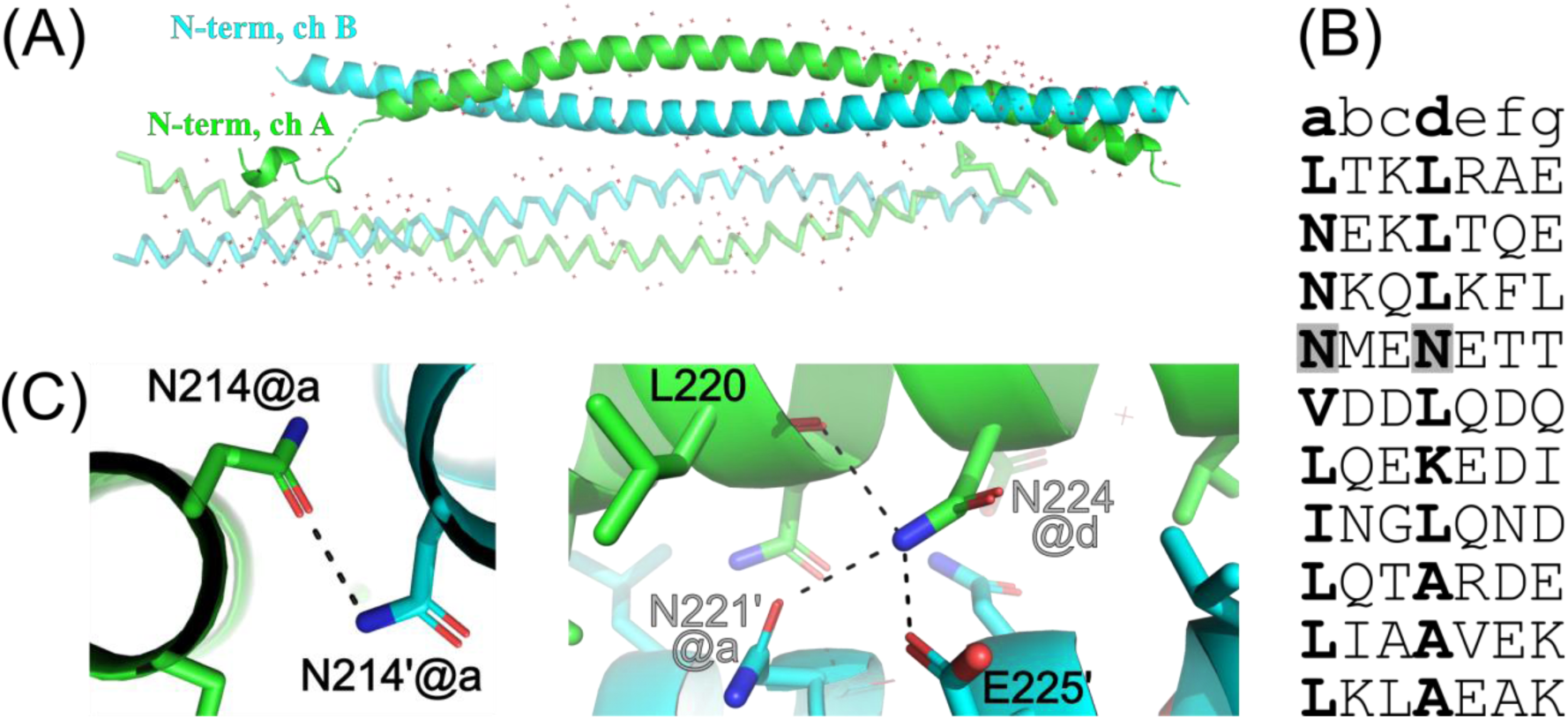
Crystal structure of the BUG1 coiled-coil domain. (A) Parallel, in-register BUG1cc dimer, with chains A and B shown in green and cyan, respectively. Symmetry-related molecules are shown in lighter shades, illustrating crystal-packing contacts involving the terminal regions. (B) Sequence of the ten heptad repeats (*a*-*g*) present in the coiled-coil structure ranging from residues 200-270 of Bug1. Core amino acid residues (*a*,*d*) are highlighted in bold and N221 and N224 are highlighted in gray boxes. (C) The left panel shows cognate N214 residues and their hydrogen-bond interaction. The right panel highlights the local environment of N224 at the coiled-coil core, showing hydrogen-bonding interactions with residues from the opposing chain, including N221′ and E225′, as well as the nearby L220 residue.

### BUG1cc forms a stable solution dimer with kinetically controlled thermal hysteresis

Having established by X-ray crystallography that BUG1cc adopts a parallel, in-register dimeric CC architecture, we next examined whether this organisation is preserved in solution and how it responds to changes in environmental conditions. Far-UV CD spectroscopy revealed the characteristic spectrum of a predominantly α-helical protein, with a positive band near 190 nm and negative bands between 208 and 222 nm (Figure 2A). Spectral deconvolution estimated 77.9% regular α-helix and 22.1% distorted α-helix, consistent with the highly helical structure observed in the crystallographic model.

**Figure 2.**
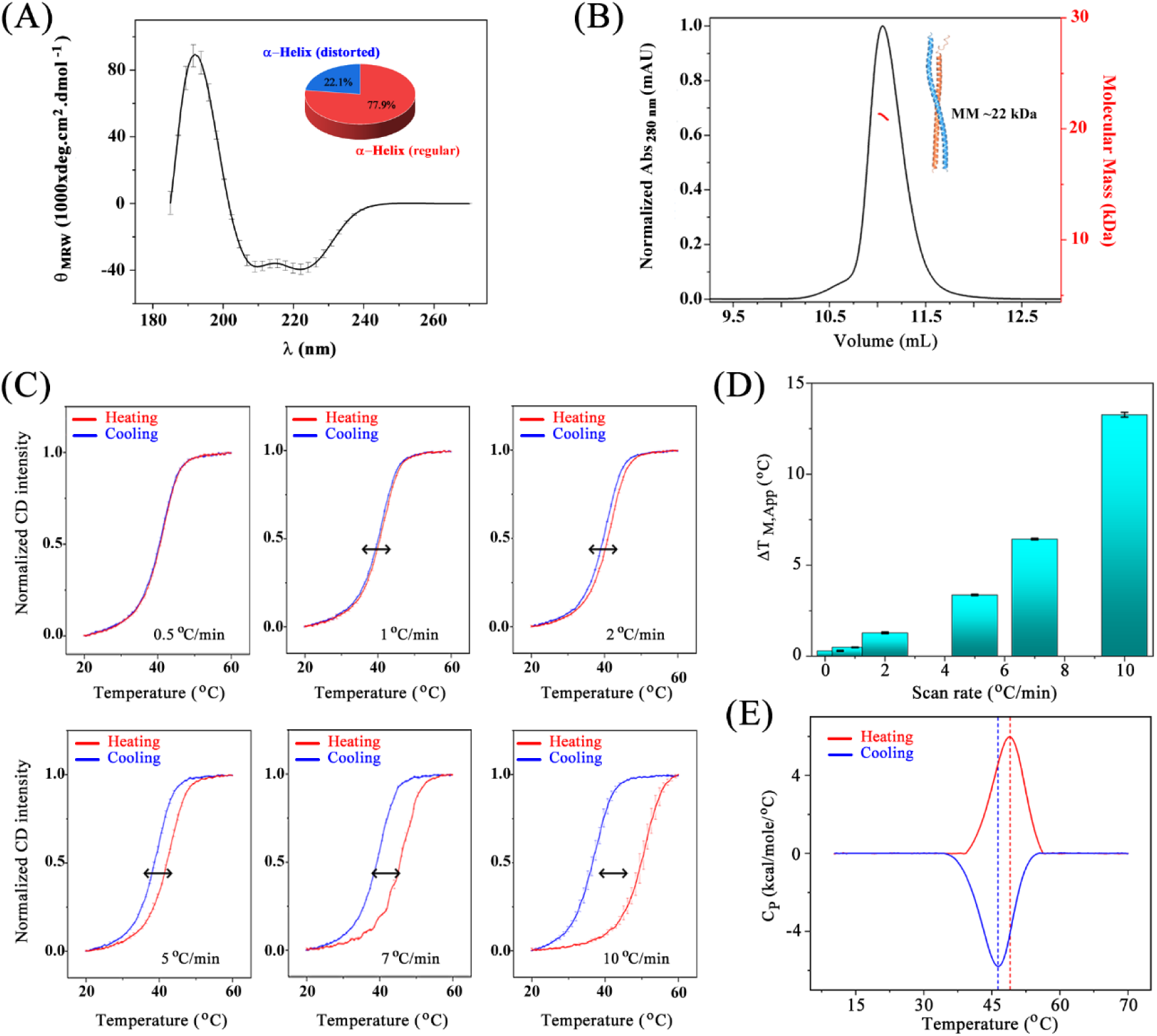
Solution-state organisation and scan-rate-dependent thermal hysteresis of BUG1cc. (A) Far-UV circular dichroism spectrum of BUG1cc, showing the characteristic signature of a predominantly α-helical protein. Spectral deconvolution indicated 77.9% regular α-helix and 22.1% distorted α-helix. (B) Size-exclusion chromatography coupled to molecular-mass determination. BUG1cc eluted as a single major species with a weight-averaged molar mass of approximately 22 kDa, consistent with a dimer in solution. (C) Thermal unfolding and refolding of BUG1cc monitored by CD at 222 nm at scan rates of 0.5, 1, 2, 5, 7, and 10 °C/min. Heating curves are shown in red and cooling curves in blue. The separation between the two branches increases progressively with scan rate, revealing pronounced thermal hysteresis. (D) Dependence of the difference between the apparent transition midpoints, ΔT_M_, on scan rate. Hysteresis is nearly negligible at the slowest scan rates and increases to approximately 13°C at 10°C/min. (E) Differential scanning calorimetry of BUG1cc at 90°C/hour. Heating and cooling thermograms are shown in red and blue, respectively, with dashed vertical lines marking the corresponding apparent transition midpoints. The calorimetric transition also displays hysteresis, with T_M_ = (48.45±0.01) °C during heating and (45.28±0.01) °C during cooling. Error bars represent the standard deviation from 3 independent measurements for the CD data. SEC-MALS and DSC were performed in 2 independent measurements.

In parallel, size-exclusion chromatography coupled to molecular-mass determination yielded a weight-averaged molar mass of approximately 22 kDa (Figure 2B), supporting a dimeric species in solution. Moreover, CD spectra collected between pH 5.5 and 8.0 retained the two negative bands near 208 and 222 nm, indicating preservation of a predominantly α-helical structure throughout this range (Figure S1A). The changes in spectral shape and the θ_222_/θ_208_ ratio suggest pH-dependent alterations in the coiled-coil environment or in helical organisation, without extensive loss of α-helical secondary structure. By contrast, increasing the NaCl concentration from below 10 mM to 150 mM produced only minor spectral changes, indicating that the α-helical structure of BUG1cc is comparatively insensitive to ionic strength under the conditions tested (Figure S1B). Together, these results support the dimeric coiled-coil observed crystallographically as the predominant solution-state architecture of BUG1cc. This organisation remains predominantly α-helical across the conditions tested, although its local helical packing appears to be modulated by pH.

We next investigated the thermal response of the dimeric BUG1cc by monitoring the CD signal at 222 nm during heating and cooling cycles performed at different scan rates (Figure 2C). At the lowest scan rate tested, 0.5 °C/min, the two signals were nearly superimposable, indicating substantial recovery of the α-helical structure when sufficient time was available for conformational relaxation. Increasing the scan rate, however, produced a progressively larger separation between the heating and cooling transitions (Figure 2C). The magnitude of this hysteresis, quantified as the difference between their apparent transition midpoints, increased from nearly negligible values at low scan rates to approximately 13 °C at 10 °C/min (Figure 2D).

The near convergence of the curves at the slowest scan rate shows that BUG1cc recovers its α-helical structure when sufficient time is available. The progressively larger separation between heating and cooling at faster ramps supports a kinetic delay in structural recovery. Thermal hysteresis arises when the timescale of the experimental perturbation becomes comparable to, or shorter than, the relaxation times of the molecular transition. Under these conditions, the conformational population lags behind the continuously changing equilibrium distribution: the structured state persists to higher temperatures during heating, whereas recovery of that state is delayed to lower temperatures during cooling. Accordingly, the transition midpoints measured in the two directions are apparent, scan-rate-dependent quantities rather than a unique equilibrium T_M_.

Thermal hysteresis measurements have, for example, been used to map the folding and assembly energy landscapes of multimeric biomolecular systems, in which the displacement of the heating and cooling branches reflects the finite rates of structural disassembly and reassembly (*35*). Scan-rate-dependent thermal shifts have similarly been used to characterise kinetically controlled transitions and estimate activation barriers in oligomeric helical systems, including collagen triple helices (*36*). For BUG1cc, structural recovery is likely kinetically demanding because it requires not only restoration of α-helicity but also productive reassociation of the two chains. Coiled-coil folding and oligomerisation are intrinsically coupled, as the same core residues contribute to helix stabilisation, inter-chain packing, orientation, oligomeric state, and register (*18*). Recovery may therefore require productive molecular encounter, conformational rearrangement of the individual chains, and correct re-establishment of the inter-chain interface (*37*).

Differential scanning calorimetry provided an independent calorimetric assessment of the thermal asymmetry observed by CD. At a scan rate of 90 °C/hour, the heating and cooling thermograms displayed distinct transition midpoints, with apparent T_M_ values of (48.45±0.01) °C and (45.28±0.01) °C, respectively, corresponding to a thermal hysteresis of >3 °C (Figure 2E and Table 2). Thus, the separation between the forward and reverse transitions is not restricted to the CD signal but is also detected through the heat absorbed and released during thermal cycling.

**Table 2.** Apparent thermal and enthalpic parameters obtained from DSC heating and cooling scans of BUG1cc.

| | $T_M$ (°C) | $\Delta H_{cal}$ (kcal mol<sup>-1</sup>) | $\Delta H_{vH}$ (kcal mol<sup>-1</sup>) | $\Delta H_{vH}/\Delta H_{cal}$ |
| --- | --- | --- | --- | --- |
| **Heating** | 48.45 ± 0.01 | 49.4 ± 0.2 | 95.9 ± 0.4 | 1.94 ± 0.01 |
| **Cooling** | 45.28 ± 0.01 | -64.7 ± 0.2 | -86.0 ± 0.3 | 1.33 ± 0.01 |

\* Error bars represent standard deviation from 2 independent measurements

The calorimetric parameters also differed between the two scan directions. During heating, the calorimetric and van’t Hoff enthalpies were (49.4±0.2) kcal/mol and (95.9±0.4) kcal/mol, respectively, yielding a ΔH_vH_/ΔH_cal_ ratio of 1.94±0.01 (Table 2). With ΔH_cal_ expressed per mole of chain, the ratio near two is consistent with a cooperative transition involving the dimer, whose organisation was independently established by crystallography and SEC-MALS. However, because these parameters were obtained under conditions exhibiting thermal hysteresis, they do not establish an equilibrium two-state mechanism or resolve whether dimer dissociation and loss of helicity occur simultaneously.

During cooling, ΔH_cal_ and ΔH_vH_ were (64.7±0.2) kcal/mol and (86.0±0.3) kcal/mol, respectively, corresponding to a ratio of 1.33±0.01 (Table 2). The decrease in the apparent cooperativity ratio during cooling further indicates that the reverse transition does not retrace the heating pathway under the experimental conditions. Instead, the two branches likely sample different non-equilibrium conformational populations, consistent with the kinetic asymmetry inferred from the CD measurements.

Taken together, the DSC data support the conclusion that BUG1cc undergoes a cooperative but non-ideal thermal transition whose apparent thermodynamic parameters differ between scan directions at the applied scan rate. The ratio near two during heating is compatible with the protein’s dimeric nature, whereas the divergence between heating and cooling reinforces that the transition cannot be interpreted as a fully reversible equilibrium two-state process. These findings motivated a molecular investigation of the relationship between dimer dissociation and loss of intra-chain helicity. We therefore used structure-based simulations to determine how disruption of the coiled-coil interface relates to unfolding of the individual chains.

### Structure-based simulations reveal a single cooperative transition and a conformational barrier to reassociation

To address this question, we used the experimentally determined BUG1cc structure as the reference state for structure-based simulations, allowing us to monitor inter-chain association and intra-chain folding independently. Tracking Q_inter_ and Q_intra_ as separate coordinates shows that the two processes are not resolvable in temperature (Figure 3A). Referred to their own baselines, the helical coordinate reaches its midpoint at T* = 0.974, and the interface coordinate at T* = 0.996, a separation of 0.022 in reduced units that maps to approximately 1 °C, far below the 7.4 °C width of the calorimetric transition. The two coordinates therefore melt as a single cooperative unit, in agreement with the calorimetric ratio ΔH_vH_/ΔH_cal_ = 1.94 obtained on heating, which suggest the dimer rather than the individual chain as the cooperative unit. The chain-separation coordinate behaves as expected for a consequence rather than a third transition: d_COM_ begins to rise across the same interval and continues to increase above it as the released chains diffuse apart, so its apparent midpoint lies higher (T* = 1.063) and is not a thermodynamic transition temperature (Figure 3B).

**Figure 3.**
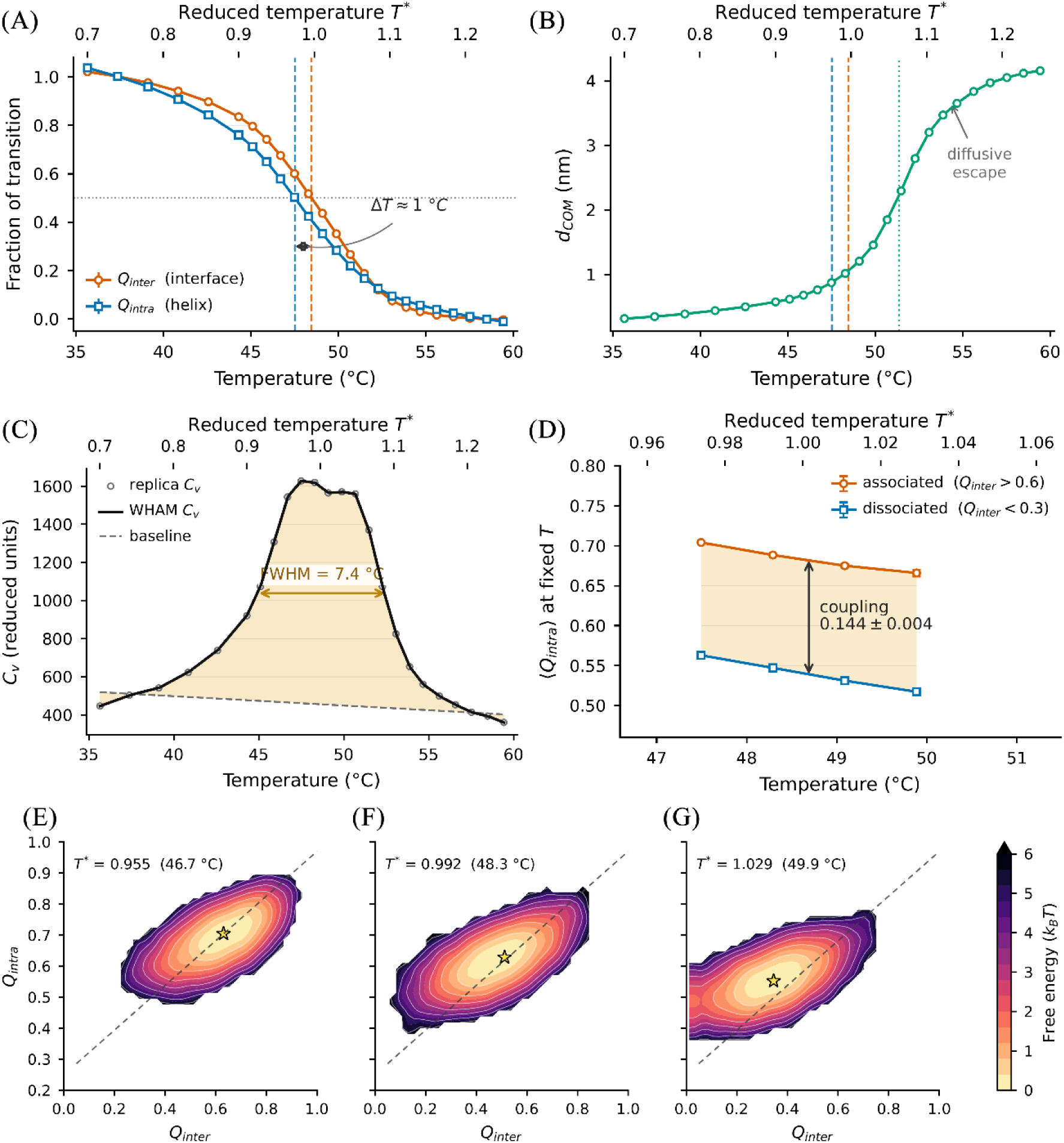
Dimer thermodynamics from REMD/WHAM. (A) Q_inter_ (interface) and Q_intra_ (helix) versus temperature, each referenced to its own pre- and post-transition baselines; the two midpoints (T* = 0.996 and 0.974; 48.45 and 47.5 °C) lie about 1 °C apart, within the transition width. (B) Inter-chain centre-of-mass distance dCOM versus temperature; separation begins across the same interval and continues above it, as expected for diffusive escape following contact loss. (C) WHAM heat capacity Cv: a single peak whose full width at half maximum (7.4 °C, measured on the baseline-subtracted profile) equals the DSC Δ*T*_1/2_ by calibration; the per-replica values and the linear baseline used to define the excess are shown. A single peak is obtained throughout. (D) Interface–helix coupling measured at fixed temperature: 〈Qintra〉 for associated (Qinter > 0.6) and dissociated (Qinter < 0.3) frames of the same replica. Losing the partner costs the chain 0.144 ± 0.004 of its native intra-chain contacts, constant across the transition region. Error bars are standard errors of the conditional means. (E–G) Free-energy surfaces F(Q_inter_, Q_intra_) at 46.7 °C (T* = 0.955, E), 48.3 °C (T* = 0.992, F) and 49.9 °C (T* = 1.029, G), from smoothed histograms of the corresponding replicas; the dashed line marks the diagonal. The minimum migrates along it, the signature of coupled rather than sequential loss of the two types of structure.

The heat-capacity profile is consistent with this picture (Figure 3C). The WHAM Cv shows a single peak, and the measured endotherm is likewise close to ideal: ΔT_1/2_ = 7.44 °C on heating, compared with 7.22 °C expected if the cooperative unit were the dimer (ΔH_vH_ ≈ 2ΔH_cal_). The simulated width is not an independent prediction because the temperature scale was calibrated on it; the model adds the reason for the width. Conditioning the ensemble on association state at fixed temperature shows that a chain inside an associated dimer retains 0.144 ± 0.004 more native intra-chain contacts than a dissociated chain, a value constant across the transition region (Figure 3D). Roughly one seventh of the helical structure of each chain is therefore held in place by the partner and is lost the moment the interface breaks. This is why no dissociated-but-folded species accumulates: there is no temperature window in which the interface can be absent while the helix remains intact, and the two coordinates are forced to melt together.

The two-dimensional free-energy surfaces provide the structural counterpart (Figure 3E– G). As temperature increases, the populated basin migrates along the diagonal of the (Q_inter_, Q_intra_) plane rather than along either axis in turn, which is the landscape signature of coupled loss of interface and helix. At the transition temperature the ensemble is broad along that diagonal, so partially dissociated and partially helical configurations are visited continuously, but they do not form a separate free-energy minimum. When mapped onto the experimental temperature scale, the model therefore reproduces the single-peaked, near-two-state appearance of the DSC endotherm while still resolving, at the level of individual contacts, the coupling that produces it. It further predicts the thermal behaviour of the isolated monomer (Figure S2), while contact-frequency maps provide a residue-level view of the same process, showing that the interface empties progressively from the chain termini inward while the local helical contacts of each chain are only partially lost (Figure S3).

Because each chain loses part of its helical structure at the moment the interface breaks, the species that must re-associate on cooling is not an unfolded random coil but a compact, partially helical monomer: across the transition region, the dissociated chains retain roughly a quarter of their native helical contacts above the denatured baseline. The WHAM free-energy landscape of the isolated monomer at its own T_M_ is a single broad, shallow basin rather than a set of discrete states (Figure 4A), with its minimum at R_g_ = 2.3 nm and R_ee_ = 5.3 nm and a mean R_ee_ of 6.5 nm, less than half the native value of 13.2 nm. The released chain is therefore collapsed and self-contacted, and conformationally heterogeneous rather than trapped in one well-defined state. Recovery of the native dimer from these compact states requires chain extension and reorganisation of intramolecular contacts as the inter-chain interface forms. This conformational requirement provides a molecular explanation for the delayed structural recovery observed in the CD experiments.

**Figure 4.**
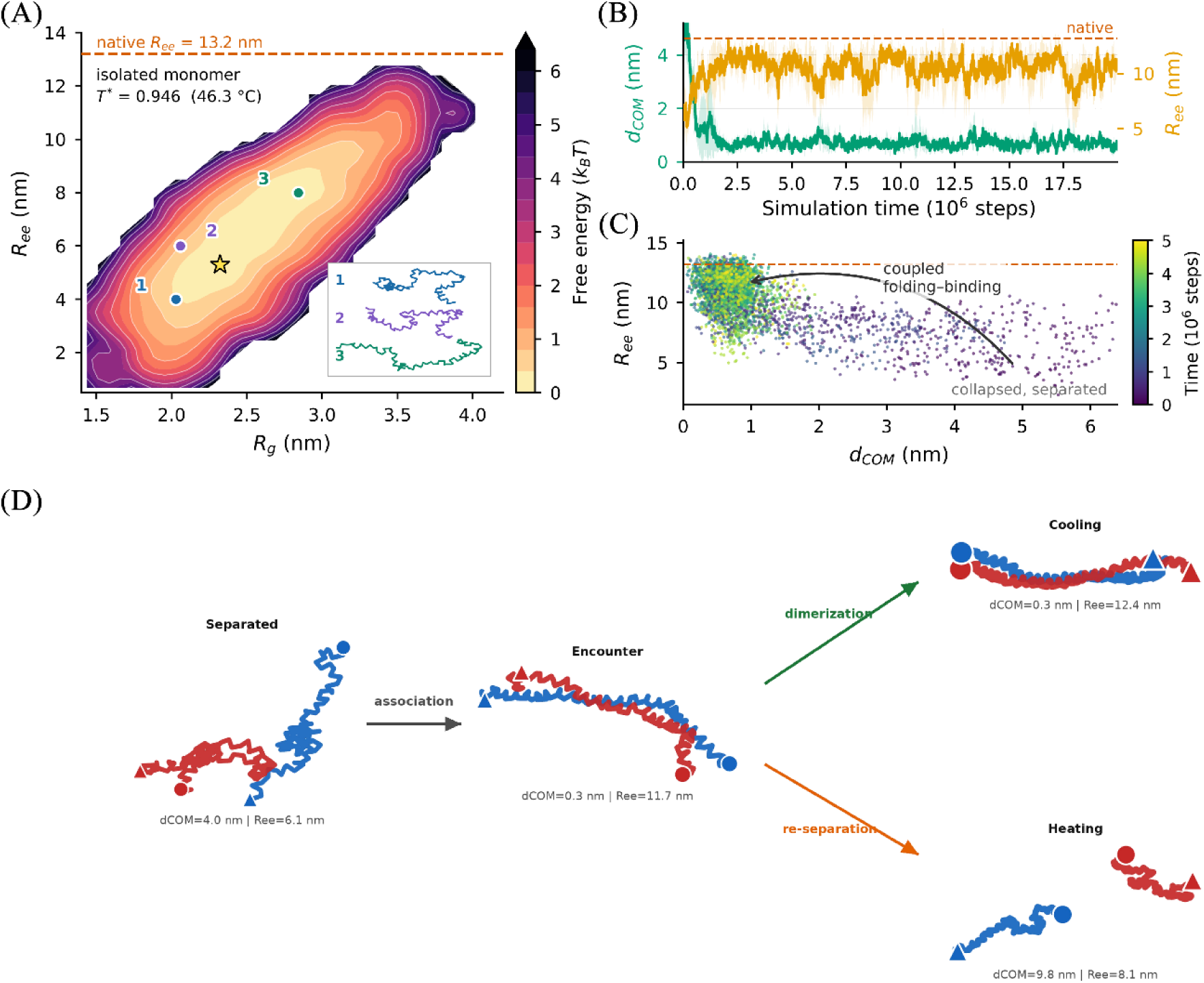
Monomer landscape and coupled folding–binding. (A) Monomer WHAM free-energy landscape of the isolated chain at T* = 0.946 (46.3 °C), the replica closest to the monomer TM, over (Rg, Ree), contoured in kBT: a single broad, shallow basin rather than a set of well-separated states, with its minimum at Rg = 2.3 nm and Ree = 5.3 nm and a mean Ree of 6.5 nm; the native Ree = 13.2 nm (dashed line) lies far outside the populated region. The inset shows representative Cα conformations sampled at Ree = 8, 6 and 4 nm. (B) Encounter time series (20 M steps; mean ± s.d. over the four converged replicates): mean dCOM collapses from ≈ 4 nm to ≈ 0.7 nm within ≈ 2 M steps, concurrent with a rise in mean Ree toward the native reference (13.2 nm), so the encounter drives chain extension rather than preceding it. (C) Encounter phase space (dCOM vs Ree, coloured by time): the productive route runs along the coupled folding–binding diagonal from separated/collapsed to extended/dimerised. (D) Structural pathway with measured coordinates: separated (dCOM = 4.0 nm, Ree = 6.1 nm) → encounter (dCOM = 0.3 nm, Ree = 11.7 nm) → cooling: stabilised dimer (dCOM = 0.3 nm, Ree = 12.4 nm) / heating: re-separated monomers (dCOM = 9.8 nm, Ree = 8.1 nm).

The encounter simulations show that folding and binding are coupled rather than sequential: as two monomers in compact separated states approach one another, each chain extends towards the native R_ee_ of 13.2 nm while d_COM_ decreases, rather than completing extension before contact (Figure 4B). The interface therefore forms while the chain is still extending, and the productive dimerisation route traces a coupled folding–binding diagonal in the (d_COM_, R_ee_) phase space from the separated/collapsed corner to the extended/dimerised corner (Figure 4C). Figure 4D summarises the overall pathway: separated monomers form a partially extended encounter complex at the dimer T_m_ that either locks into a stabilised dimer on cooling or re-separates on heating. Driving the shared encounter state via cooling or heating branches reproduces the directional asymmetry that underlies hysteresis (Figure S4). The fixed-temperature all-atom simulations are consistent with this picture in a complementary sense: over 100 ns at 25– 60 °C the dimer remains intact and fully helical at every temperature (Q > 0.98, helicity 0.92– 0.95, R_g_ and SASA invariant), confirming that neither dissociation nor helix melting is accessible to unbiased all-atom sampling on this timescale and that the coarse-grained REMD approach is required to reach the thermodynamics of the transition.

To examine the experimental implications of the coupling between chain association and conformational reorganisation suggested by the simulations, we measured the thermal response of BUG1cc at concentrations from 0.02 to 0.6 mg mL^-1^, using fixed heating and cooling rates of 5 °C min^-1^. Increasing protein concentration from 0.02 to 0.4 mg mL^-1^ shifted both heating and cooling transitions upwards by approximately 9 °C, with no further increase at 0.6 mg mL^-1^ (Figure 5A). Despite these substantial shifts, thermal hysteresis remained approximately 5–6 °C across the concentration range, with no systematic decrease at higher concentrations (Figure 5B). Thus, concentration strongly affected the location of the dominant transitions while having comparatively little effect on their separation under the applied scan conditions. The concentration dependence of the apparent transition temperatures supports chain association in the thermal response, while the persistence of hysteresis suggests that conformational rearrangements also contribute to delayed structural recovery, consistent with the mechanism proposed by the simulations.

**Figure 5.**
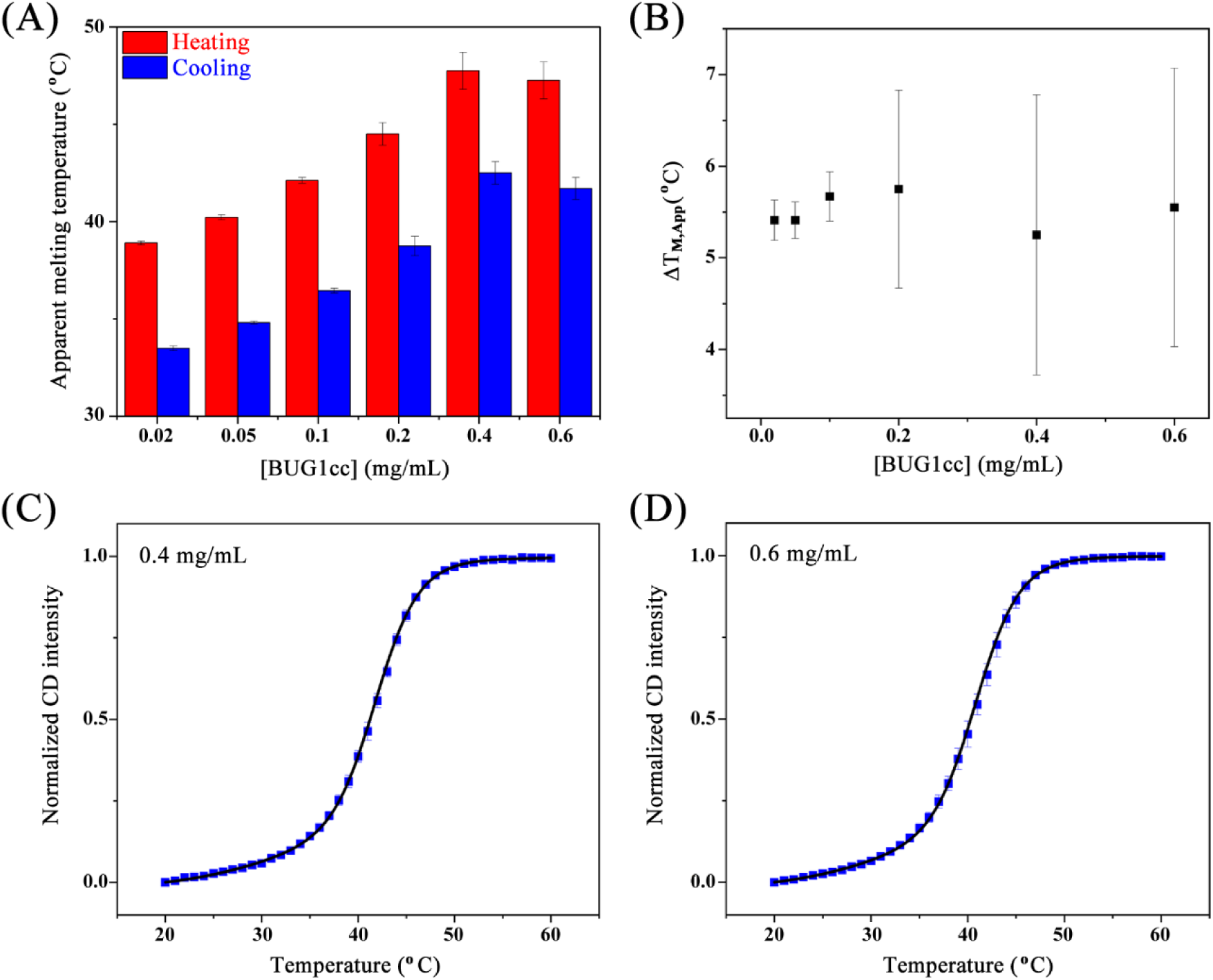
Concentration dependence of the thermal response and hysteresis of BUG1cc. (A) Apparent transition temperatures determined from the dominant peak in the magnitude of the first derivative of the CD signal at 222 nm during heating (red) and cooling (blue), at protein concentrations of 0.02–0.6 mg mL^-1^. All scans were performed at 5 °C min^-1^. (B) Thermal hysteresis as a function of protein concentration. (C–D) Normalised CD cooling profiles at 0.4 and 0.6 mg mL⁻¹, respectively. Blue squares represent experimental data and black lines represent double-sigmoid fits used to describe the asymmetric profiles. Cooling proceeds from high to low temperature. The fitted components are phenomenological and do not represent independently resolved molecular transitions. Data are presented as mean ± standard deviation from four independent measurements.

The cooling profiles at 0.4 and 0.6 mg mL⁻¹ provided further insight into structural recovery. Both showed similar asymmetry, with systematic deviations from single-sigmoid fits (Figure S5) captured by double-sigmoid fits (Figure 5C–D). At both concentrations, the fits described a dominant change in ellipticity, followed during cooling by a broader contribution accounting for approximately 18–19% of the fitted signal amplitude. The similar component widths and relative amplitudes indicate a consistent pattern of structural recovery (Table 3). This broader contribution appeared as an asymmetry of the cooling profile, which retained a single dominant peak in its first derivative. Together with the concentration-dependent shifts and persistent hysteresis, this gradual completion of structural recovery supports conformational rearrangements during coiled-coil assembly, consistent with the coupled association and chain extension observed in the simulations.

**Table 3.**
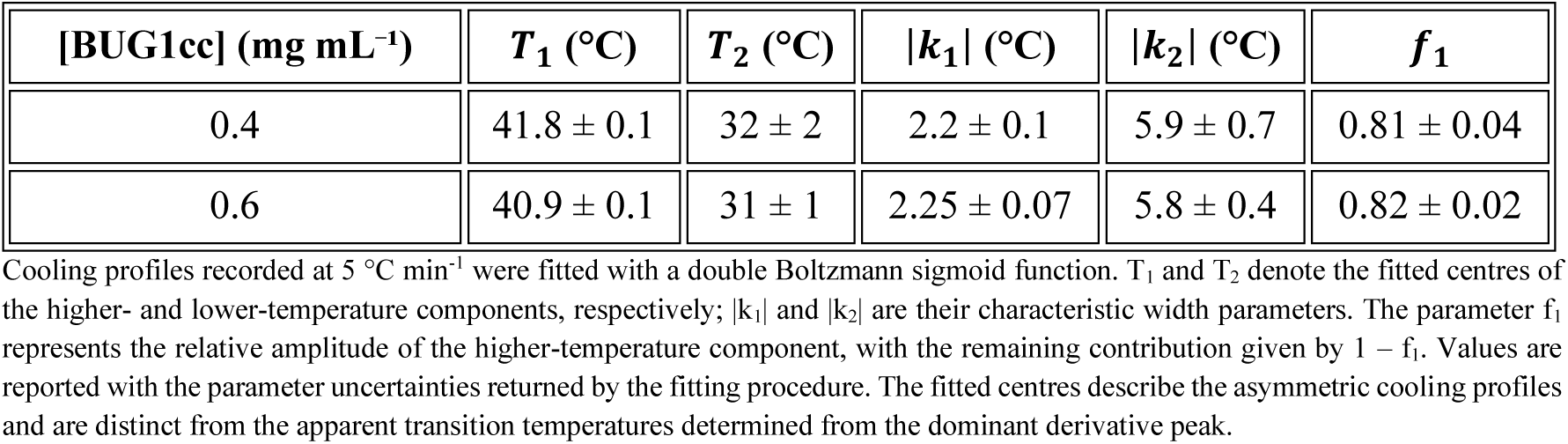
Parameters of double-sigmoid fits to BUG1cc cooling profiles.

| [BUG1cc] (mg mL<sup>-1</sup>) | $T_1$ (°C) | $T_2$ (°C) | $\vert k_1\vert $ (°C) | $\vert k_2\vert $ (°C) | $f_1$ |
| --- | --- | --- | --- | --- | --- |
| 0.4 | 41.8 ± 0.1 | 32 ± 2 | 2.2 ± 0.1 | 5.9 ± 0.7 | 0.81 ± 0.04 |
| 0.6 | 40.9 ± 0.1 | 31 ± 1 | 2.25 ± 0.07 | 5.8 ± 0.4 | 0.82 ± 0.02 |

Cooling profiles recorded at 5 °C min<sup>-1</sup> were fitted with a double Boltzmann sigmoid function. $T_1$ and $T_2$ denote the fitted centres of the higher- and lower-temperature components, respectively; $|k_1|$ and $|k_2|$ are their characteristic width parameters. The parameter $f_1$ represents the relative amplitude of the higher-temperature component, with the remaining contribution given by $1 - f_1$ . Values are reported with the parameter uncertainties returned by the fitting procedure. The fitted centres describe the asymmetric cooling profiles and are distinct from the apparent transition temperatures determined from the dominant derivative peak.

The concentration series helps constrain the possible origins of thermal hysteresis. Increasing chain concentration twenty-fold, from 0.02 to 0.4 mg mL^-1^, shifted both apparent transition temperatures upwards by approximately 9 °C, while their separation remained approximately 5–6 °C (Figure 5A–B). These observations support an association-dependent thermal response and suggest that increasing the frequency of intermolecular encounters is insufficient to eliminate the delay in structural recovery under the conditions tested. The simulations provide a plausible molecular explanation: dissociated chains populate compact, partially helical conformations and undergo extension and reorganisation as the native interface forms. The similar widths and relative amplitudes of the fitted cooling components at 0.4 and 0.6 mg mL⁻¹ are consistent with a persistent pattern of structural recovery (Table 3), although these phenomenological components do not identify individual kinetic steps. Together, the results support a contribution from conformational rearrangements to delayed reassociation.

## Conclusion

This study establishes the atomic architecture of a predicted coiled-coil region of the yeast Golgi-associated protein Bug1. The crystal structure revealed a parallel, in-register homodimer stabilised by a canonical heptad-repeat arrangement and specific polar interactions within the coiled-coil core. Consistently, SEC-MALS established the dimeric organisation of BUG1cc in solution, while CD showed that its highly α-helical structure is preserved across the pH and ionic strength conditions examined. Together, these results support the coiled-coil architecture identified crystallographically as a robust solution state.

The thermal measurements connect this architecture to the folding and assembly behaviour of BUG1cc. CD revealed scan-rate-dependent hysteresis, independently corroborated by DSC. Increasing protein concentration shifted the apparent heating and cooling temperatures while preserving approximately 5-6 °C of hysteresis, supporting an association-dependent thermal response in which conformational rearrangements contribute to delayed recovery. The asymmetric cooling profiles at higher concentrations further support a gradual completion of structural recovery. Structure-based simulations provide a molecular framework for these observations, showing that the coiled-coil interface and the helicity of each chain are thermodynamically coupled and melt as a single cooperative unit, and that the monomer released on dissociation is compact and only partially helical, so that reassociation couples chain extension to interface formation and is limited by a conformational rather than a bimolecular step.

These results establish BUG1cc as an experimentally validated dimeric coiled-coil and connect its molecular architecture to a cooperative thermal response with asymmetric recovery kinetics. More broadly, the study reinforces the value of direct structural and biophysical characterisation of golgin coiled-coil regions and provides a framework for investigating how chain association and conformational reorganisation jointly shape their assembly dynamics.

## Acknowledgements

The authors thank the Brazilian funding agencies CAPES (Coordination for the Improvement of Higher Education Personnel), FAPESP (Fundação de Amparo à Pesquisa do Estado de São Paulo; grants #2022/06006-0, #2023/12516-3, #2023/13459-3, #2023/01744-5, and #2025/19163-4), and CNPq/MCTI (grant #404982/2025-5) for financial support. We also thank the Brazilian Center for Research in Energy and Materials (CNPEM) and the MANACÁ beamline staff for assistance during the X-ray diffraction data collection experiments (proposal #20242586). Finally, we thank the National Laboratory for Scientific Computing (LNCC/MCTI, Brazil) for providing high-performance computing (HPC) resources on the SDumont supercomputer (project prodex-GRB2; http://sdumont.lncc.br).

## Declaration of generative AI and AI-assisted technologies in the writing process

While preparing this work, the authors used ChatGPT-5 and Grammarly AI Writing Assistance solely to improve the language and readability of the text. After using these tools/services, the authors reviewed and edited the content as needed and take full responsibility for the publication’s content.s

