## Supplementary Information for "Experimental Validation of Coiled-Coil Architecture and Folding Dynamics in the Golgin Bug1"

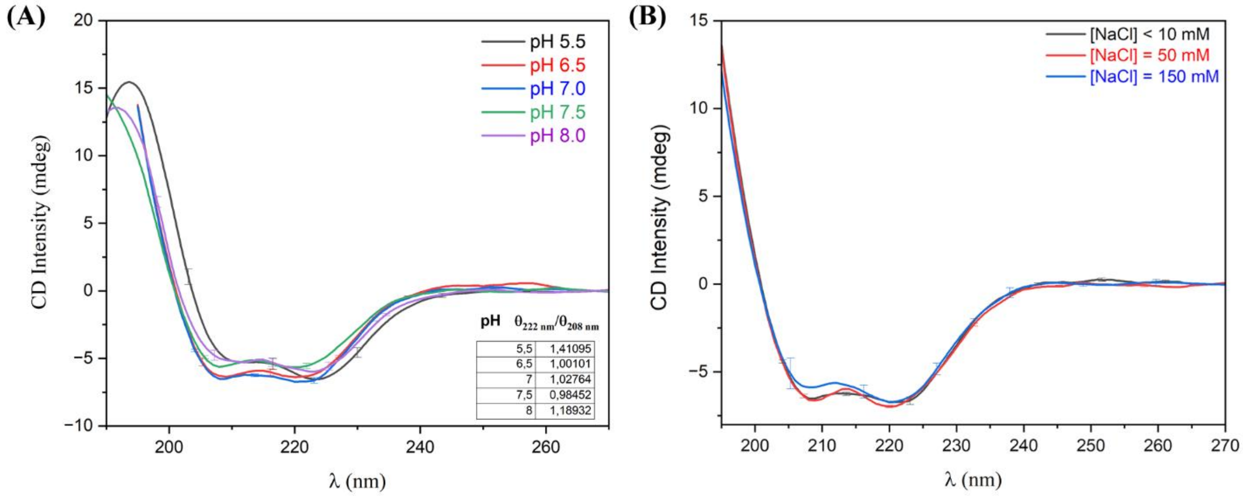


**Figure S1**. Effect of pH and ionic strength on the secondary structure of the Bug1 coiled-coil domain at 0,1 mg/mL fixed concentration. Far-UV circular dichroism (CD) spectra recorded (A) at pH values ranging from 5.5 to 8.0 and (B) at NaCl concentrations below 10, 50, and 150 mM. Under all conditions, the spectra display the characteristic double minimum near 208 and 222 nm associated with an α-helical coiled-coil structure. The $\theta_{222}/\theta_{208}$ ratios obtained at each pH are shown in the inset of panel A. Overall, the spectral profiles indicate that the α-helical organisation of the Bug1 coiled-coil is preserved across the evaluated pH and ionic-strength ranges. Error bars represent standard deviation from 3 independent measurements.

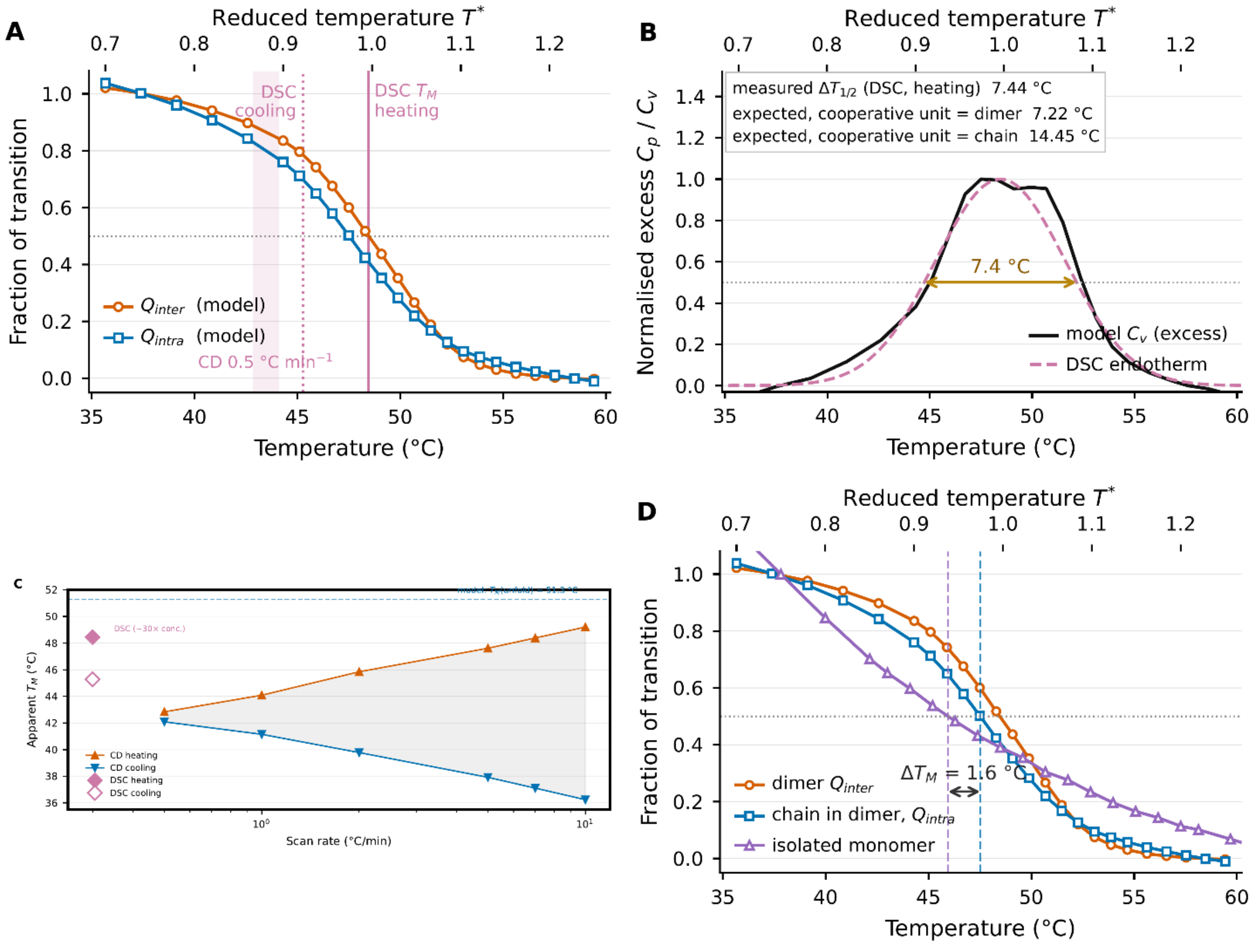


**Figure S2.** Model predictions mapped to experiment. (A) $Q_{inter}$ and $Q_{intra}$ on the experimental Celsius axis, each referred to its own pre- and post-transition baselines. The interface midpoint is anchored to the DSC heating $T_M$ (48.45 °C, by construction) and the helical midpoint falls 0.9 °C below it (47.5 °C), so the two coordinates are indistinguishable within the width of the transition. Vertical lines mark the DSC heating and cooling midpoints; the shaded band spans the CD quasi-equilibrium midpoints at 0.5 °C/min (cooling 42.84 °C, heating 44.09 °C). (B) Width of the transition: the baseline-subtracted excess $C_v$ of the model compared with a Gaussian representation of the DSC endotherm. Because the temperature scale was calibrated on the DSC $\Delta T_{1/2}$, this panel documents the calibration rather than testing it; the informative comparison is with the reference widths in the inset, the measured 7.44 °C being close to the 7.22 °C expected if the cooperative unit were the dimer and far from the 14.45 °C expected for a single chain. (C) Hysteresis reference: apparent CD midpoints on heating (dissociation-limited) and cooling (re-association-limited) diverge with increasing scan rate ($\Delta T_M$ = 1.25 °C at 0.5 °C/min → 7.0 °C at 5 °C/min → 13.1 °C at 10 °C/min); the DSC midpoints, recorded at 90 °C/h on a 30-fold more concentrated sample, sit above the CD trend, as expected for a bimolecular dissociation whose $T_M$ depends on chain concentration. (D) Falsifiable prediction: replica-exchange simulations of the isolated chain give a melting midpoint 1.6 °C below that of the same chain within the dimer, the cooperative stabilisation contributed by the partner. A monomeric variant, or measurements at sufficiently low chain concentration, should therefore melt below the dimer rather than at the same temperature.

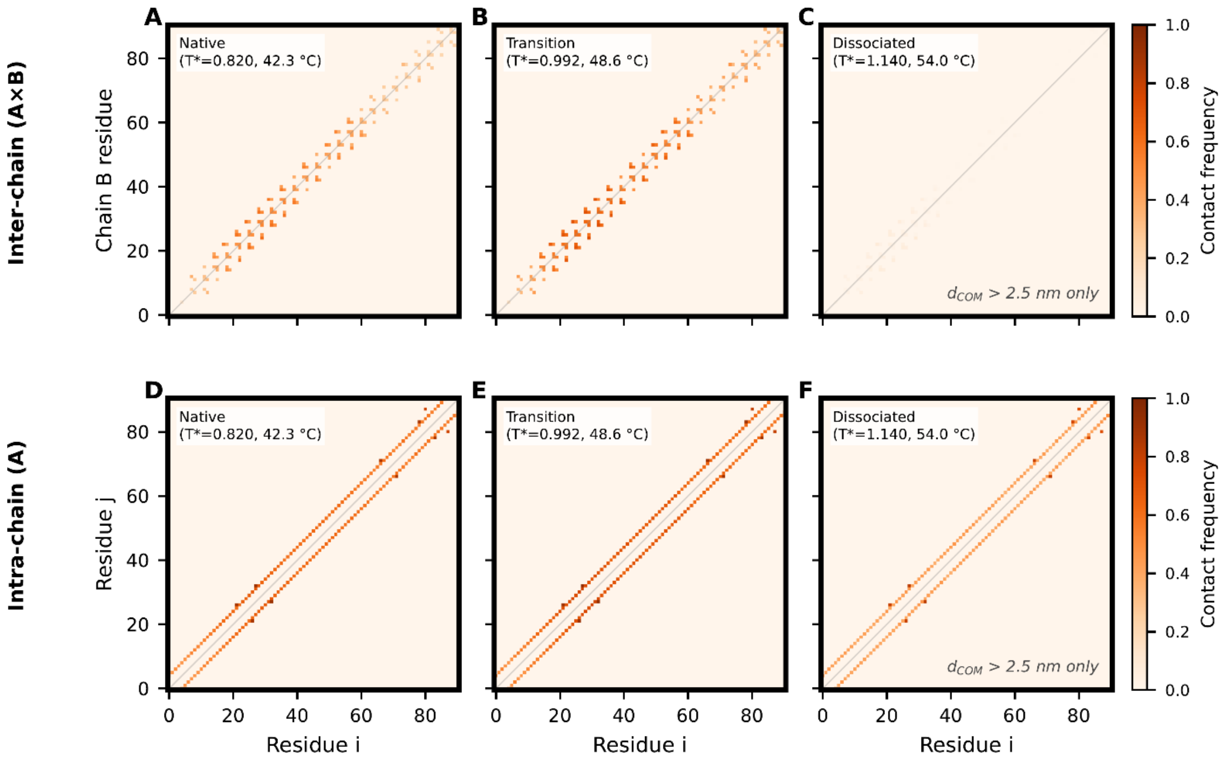


**Figure S3.** Residue × residue contact-frequency maps at three thermal states. (A–C) Inter-chain interface contacts (chain A × chain B) at T* = 0.820, 0.992 and 1.140 (42.3, 48.6 and 54.0 °C), the dissociated map being restricted to frames with $d_{COM} > 2.5$ nm: the interface empties progressively from the chain termini inward as temperature rises, with the central contacts persisting to the highest temperature. (D–F) Intra-chain contacts within chain A at the same three temperatures retain a substantial fraction of their native frequency even in the dissociated ensemble, the residue-level counterpart of the compact, partially helical monomer released on dissociation.

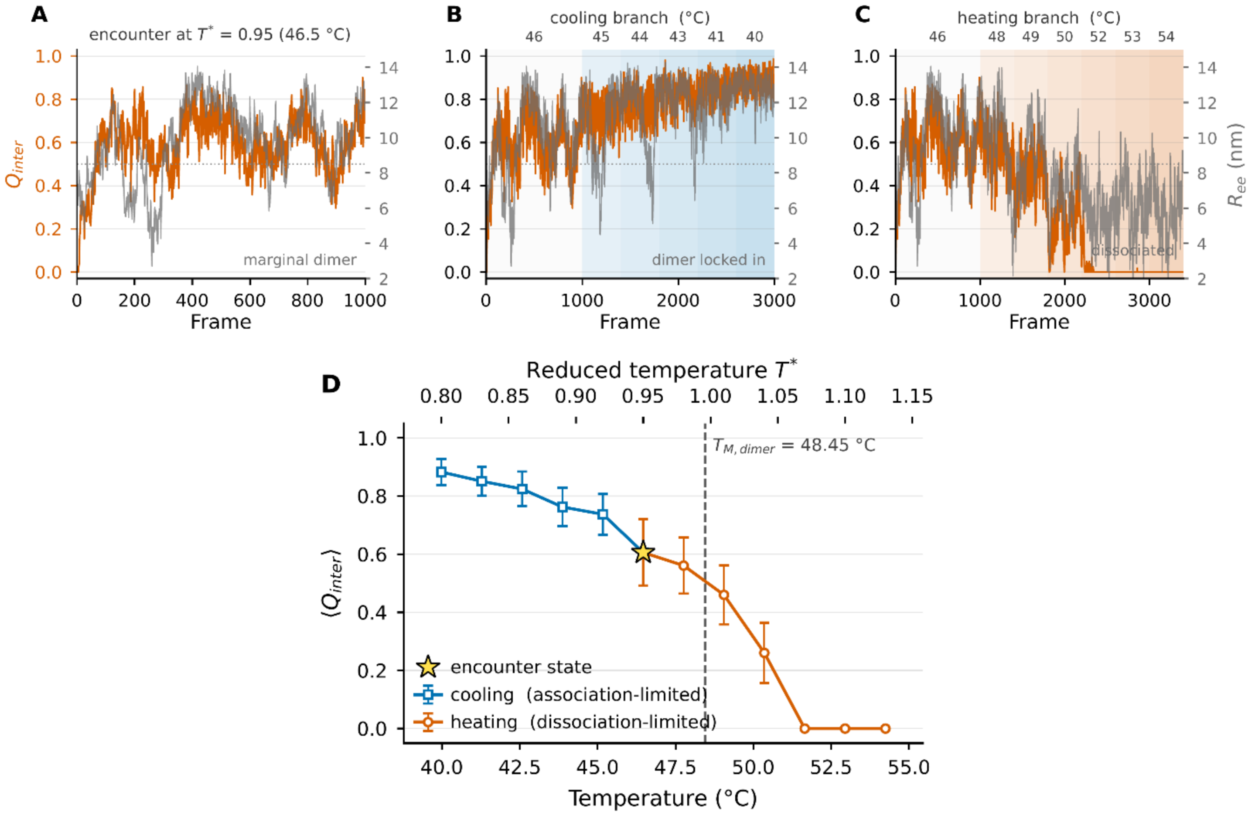


**Figure S4.** Encounter dynamics and directional asymmetry. (A) $Q_{inter}$ (orange, left axis) and $R_{ee}$ (grey, right axis) versus simulation frame during an encounter just below the dimer $T_M$ ($T^*$ = 0.95, 46.5 °C): $Q_{inter}$ fluctuates around an intermediate value, confirming a metastable encounter complex. (B) Cooling branch ($T^*$ = 0.92 → 0.80, i.e. 45.2 → 40.0 °C; shaded steps labelled in °C along the top): $Q_{inter}$ rises above 0.7 and stabilises, and $R_{ee}$ approaches the native value (native-dimer lock-in). (C) Heating branch from the same encounter state ($T^*$ = 0.98 → 1.13, i.e. 47.8 → 54.2 °C): $Q_{inter}$ decays to zero as the chains re-separate. (D) Mean $Q_{inter}$ over the last 200 frames of each step along the cooling (association-limited) and heating (dissociation-limited) branches, on a common temperature axis around $T_{M,dimer}$ = 48.45 °C; error bars are standard deviations and the star marks the shared encounter state. The two branches trace different curves, the directional asymmetry underlying scan-rate-dependent CD hysteresis, because re-association requires conformational escape from compact monomer states whereas dissociation requires only thermal weakening of the native interface.

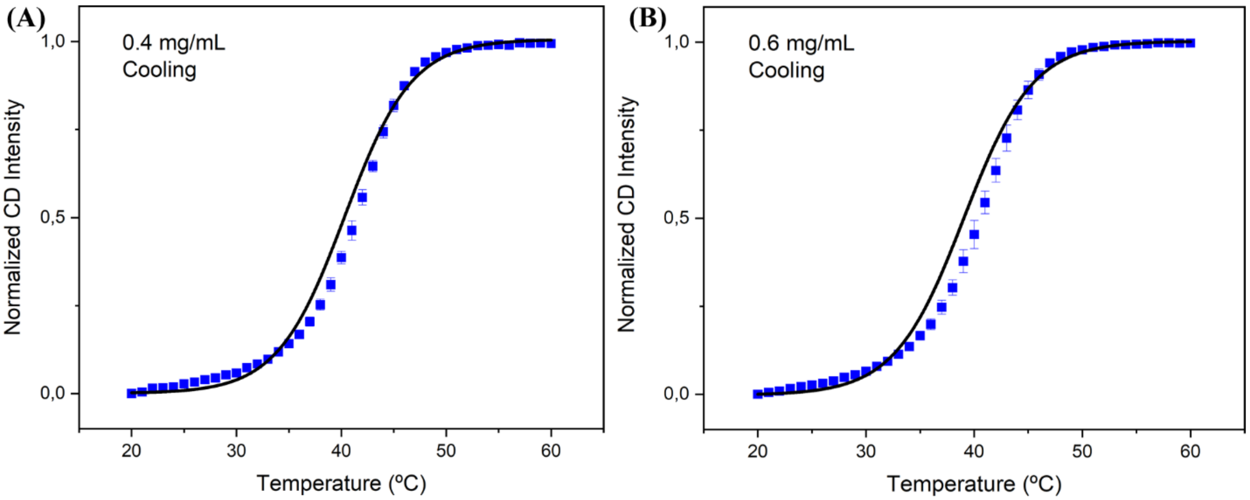


**Figure S5.** Single-sigmoid fits to BUG1cc cooling profiles at high protein concentrations. Normalised CD signals at 222 nm recorded during cooling at 5 °C $min^{-1}$ at (A) 0.4 mg $mL^{-1}$ and (B) 0.6 mg $mL^{-1}$. Blue squares represent the mean of four independent measurements, with error bars indicating standard deviation. Black lines represent single Boltzmann sigmoid fits with constant baselines. Systematic deviations between the experimental data and the fitted curves indicate that this function does not adequately capture the asymmetric cooling profiles. Cooling proceeds from high to low temperature.
